# Glycan reactive anti-HIV-1 antibodies bind the SARS-CoV-2 spike protein but do not block viral entry

**DOI:** 10.1101/2021.01.03.425141

**Authors:** Dhiraj Mannar, Karoline Leopold, Sriram Subramaniam

## Abstract

The SARS-CoV-2 spike glycoprotein is a focal point for vaccine immunogen and therapeutic antibody design, and also serves as a critical antigen in the evaluation of immune responses to COVID-19. A common feature amongst enveloped viruses such as SARS-CoV-2 and HIV-1 is the propensity for displaying host-derived glycans on entry spike proteins. Similarly displayed glycosylation motifs can serve as the basis for glyco-epitope mediated cross-reactivity by antibodies, which can have important implications on virus neutralization, antibody-dependent enhancement (ADE) of infection, and the interpretation of antibody titers in serological assays. From a panel of nine anti-HIV-1 gp120 reactive antibodies, we selected two (PGT126 and PGT128) that displayed high levels of cross-reactivity with the SARS-CoV-2 spike. We report that these antibodies are incapable of neutralizing pseudoviruses expressing SARS-CoV-2 spike proteins and are unlikely to mediate ADE via FcγRII receptor engagement. Nevertheless, ELISA and other immunoreactivity experiments demonstrate these antibodies are capable of binding the SARS-CoV-2 spike in a glycan-dependent manner. These results contribute to the growing literature surrounding SARS-CoV-2 S cross-reactivity, as we demonstrate the ability for cross-reactive antibodies to interfere in immunoassays.

## Introduction

The coronavirus disease-2019 (COVID-19) pandemic has had devastating effects on human health and economies. SARS-CoV-2 is the causative agent of COVID-19 and belongs to the coronavirus family of large, enveloped viruses that rely on a trimeric transmembrane spike (S) glycoprotein for host cell recognition and entry(1). The SARS-CoV-2 spike is a transmembrane protein consisting of an S1 domain, which contains the cell surface receptor binding domain (RBD), and S2 domain that includes the region that promotes viral fusion(2, 3). Due to its accessibility on the viral surface and its critical role in the viral life cycle, the SARS-CoV-2 spike represents a crucial antigen in host immune responses. As antibody responses to vaccination and COVID-19 infection are further investigated via mainstay serological methods(4–8), the potential for cross-reactive epitopes within SARS-CoV-2 S to provide confounding results remains an important consideration. Cross-reactive antibodies against viruses both closely and distantly related to SARS-CoV-2 may bind the spike protein with either high or low affinity and may provide the potential for virus neutralization. Indeed, several studies have demonstrated SARS-CoV-2 S cross-reactivities involving closely related coronaviruses(9– 12) along with others such as dengue virus(13, 14) and HIV(15). Investigation of such cross-reactive interactions may reveal novel viral epitopes and vulnerabilities, along with providing context for the interpretation of serological assay studies, of which some have failed to demonstrate antibody titers as an appreciable predictor of immunity, disease status, and disease progression for COVID-19(16–19).

Viral spike proteins are typically populated by a variable array of host-derived glycans, which serve multiple functions, amongst which include epitope occlusion and immune system evasion(20). Such viral glycosylation patterns themselves may be sufficiently antigenic to elicit antibody responses, and even confer viral neutralization when bound. Indeed, there exist multiple broadly neutralizing antibodies against the HIV spike gp120, whose epitopes involve critical glycan contacts(21). The SARS-CoV-2 spike contains 22 N linked glycosylation sites which are variably glycosylated(22), representing the potential for glyco-epitope mediated cross reactivity. A recent study has reported SARS-CoV-2 S glyco-epitope recognition by mannose directed Fab-dimerized antibodies against HIV gp120(15), confirming that glycosylation mediated cross-reactivity is possible. We aimed to further investigate such cross-reactivities using a panel of broadly neutralizing anti-HIV antibodies which recognize glycan and peptide epitopes within gp120.

## Results

To investigate the existence of cross-reactive glyco-epitopes within the SARS-CoV-2 spike, we subjected a panel of glycan-reactive anti-HIV-1 gp120 antibodies to an ELISA-based cross-reactivity screen (Figure 1). We selected nine anti-gp120 antibodies whose epitopes have been shown to involve glycans, along with two anti-gp120 antibodies which recognize the gp120 CD4-binding site as negative controls (Table S1) and assessed the ability of these antibodies to bind SARS-CoV-2 spike ectodomain, purified to homogeneity (Fig. S1a). While the CD4-binding site directed antibodies VRC01 and VRC03 were unable to bind the SARS-CoV-2 spike ectodomain, we observed various levels of cross-reactivity for most of the screened glycan-reactive antibodies, the most potent of which were 2G12, PGT128 and PGT126 (Figure 1). As expected, a positive control antibody V_H_-FC ab8 which targets the SARS-CoV-2 receptor-binding domain (RBD)(23) demonstrated potent binding of the SARS-CoV-2 spike ectodomain in the same assay (Figure S2a). Glycan-dependent cross-reactivity of 2G12 with the SARS-CoV-2 spike has been recently described(15), and as 2G12, PGT128 and PGT126 exhibited similar levels of cross-reactivity, we selected these antibodies for further investigation.

**Figure 1.**
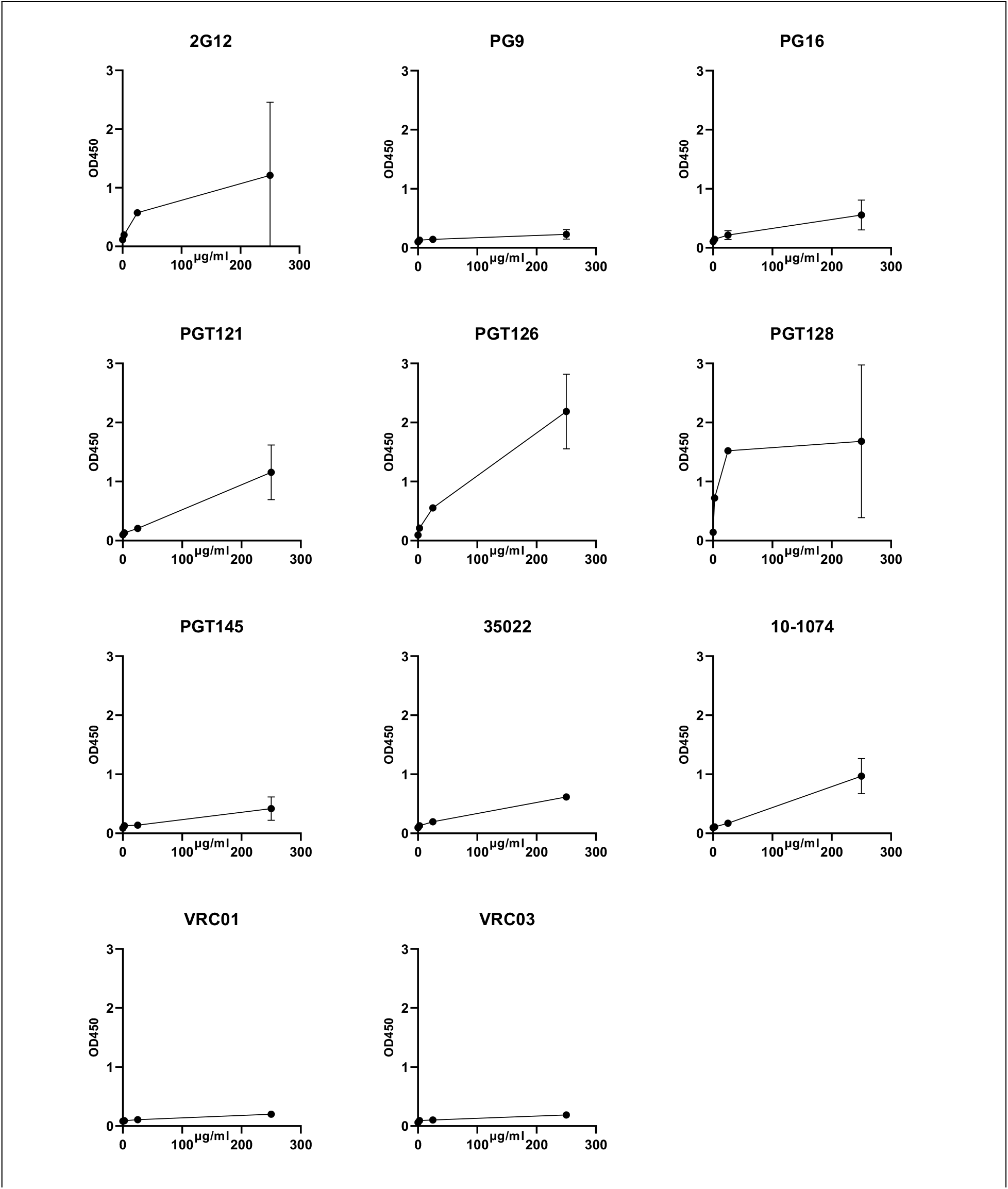
ELISA screen of glycan directed anti-gp120 antibody cross-reactivities to the SARS-CoV-2 Spike. Serial dilutions of the indicated mAbs were assessed for SARS-CoV-2 S protein binding. VRC01 and VRC03 target the CD4 binding site within gp120 and are included here as negative controls. All ELISAs were performed using BSA-based buffers (see methods). Experiments were done in duplicate; error bars indicate standard deviation (n = 2).

We next sought to determine the neutralization capabilities of these cross-reactive antibodies, via use of a SARS-CoV-2 S pseudotyped virus entry assay(24) (Figure 2). No neutralization capabilities were detected for 2G12, PGT128, and PGT126 over a wide range of concentrations, while V_H_-FC ab8 demonstrated potent neutralization, in keeping with previous reports(23). As antibody-dependent enhancement (ADE) of infection has been described for SARS-CoV-2 via engagement of cellular FcγRII receptors(25), we evaluated the potential for antibody directed enhancement of infection (ADE) by these antibodies, in the FcγRII receptor expressing K562 erythroleukemic cell line(26) (Figure 2c). We were unable to observe an enhancement of pseudoviral entry into K562 cells under a wide range of antibody concentrations, suggesting these antibodies are unlikely to mediate ADE via FcγRII receptor engagement.

**Figure 2.**
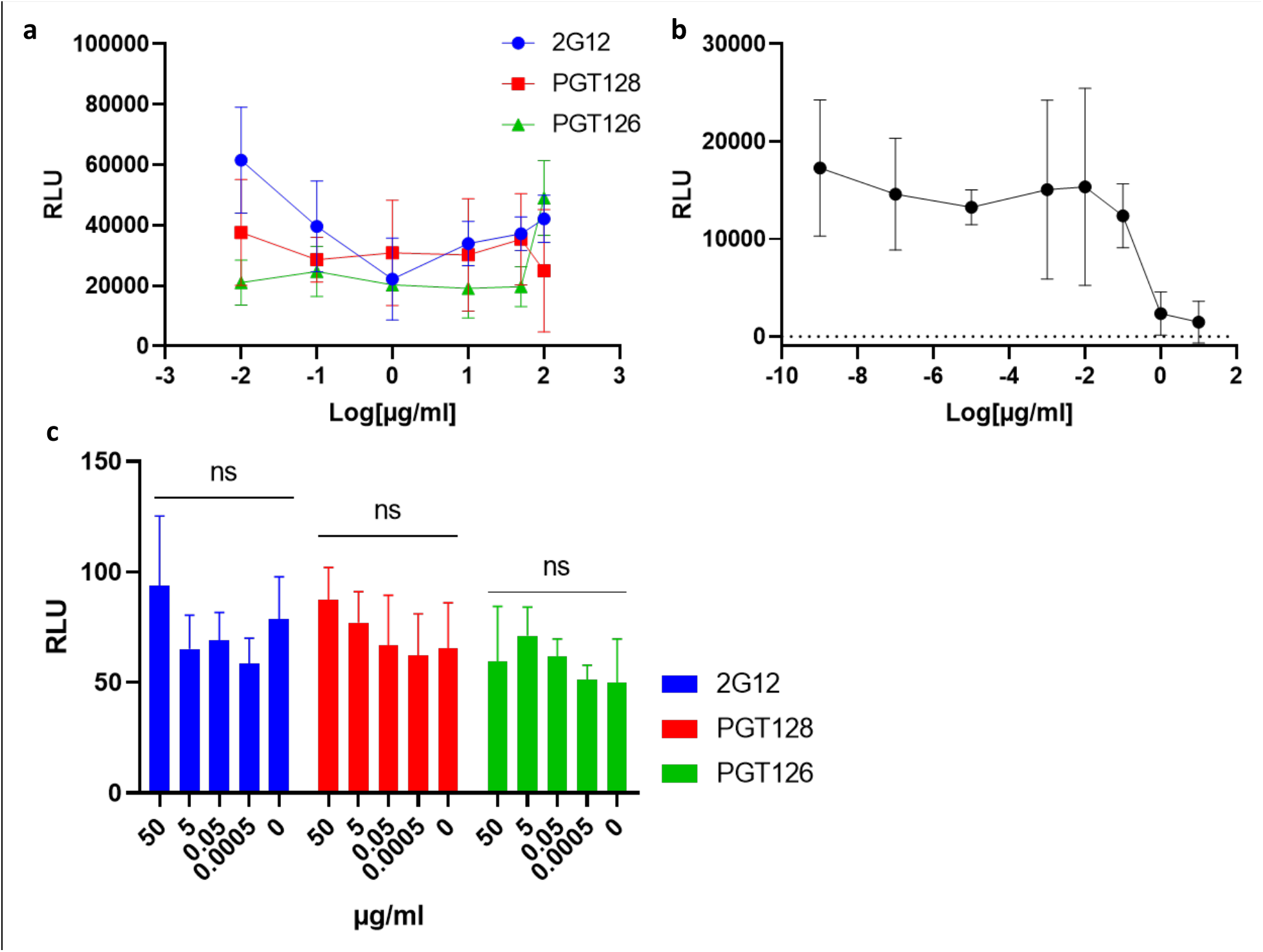
2G12, PGT128 and PGT126 do not neutralize SARS-CoV-2 S pseudo-typed virus. **(a)** HEK293-T cells stably overexpressing ACE-2 were incubated with SARS-CoV-2 S pseudo-typed virus harbouring a luciferase reporter gene, in the presence of serial dilutions of the indicated anti-gp120 antibodies. Luciferase activities in cellular lysates were determined 48 hours post-infection. (RLU: relative luciferase units). **(b)** Neutralizing antibody VH-FC ab8 subjected to the same assay described in (a). **(c)** Evaluation of antibody-dependent enhancement (ADE) for 2G12, PGT126 and PGT128 on the Fcγ RII expressing cell line: K562. SARS-CoV-2 S pseudo-type cell entry assays were performed on K562 cells as in **(a-c)**. Experiments were done in triplicate; error bars indicate standard deviation (n = 3). Statistical significance was tested by two-way ANOVA with Dunnett post-test (p > 0.05 [ns, not significant], p ≤ 0.05 [∗], p ≤ 0.01 [∗∗], p ≤ 0.001 [∗∗∗]).

To further characterize the cross-reactivity of 2G12, PGT128 and PGT126 with the SARS-CoV-2 spike protein, we assessed the immunoreactivity of these antibodies with the SARS-CoV-2 spike ectodomain via Western blot (Figure 3a). Immunoreactivity was observed for 2G12, PGT128, and PGT126, while VRC01 immunoreactivity was not detected, consistent with the results of our initial ELISA screen. We next aimed to assess the ability of these antibodies to cross-react with full-length SARS-CoV-2 spike under native conditions. To this end, we performed immunoprecipitation experiments utilizing lysate generated from cells transiently expressing the full-length SARS-CoV-2 spike (Figure 3b). The full-length SARS-CoV-2 spike was successfully immunoprecipitated by 2G12, PGT128 and PGT126, but not by VRC01, implicating these cross-reactive epitopes to be present in the full-length spike under non-denaturing conditions. Furthermore, we demonstrate dose-dependent interactions between 2G12, PGT128, and PGT126 but not VRC01 and cell-associated full-length SARS-CoV-2 spike via a cell-based ELISA assay (Figure 3c), corroborating our immunoprecipitation results.

**Figure 3.**
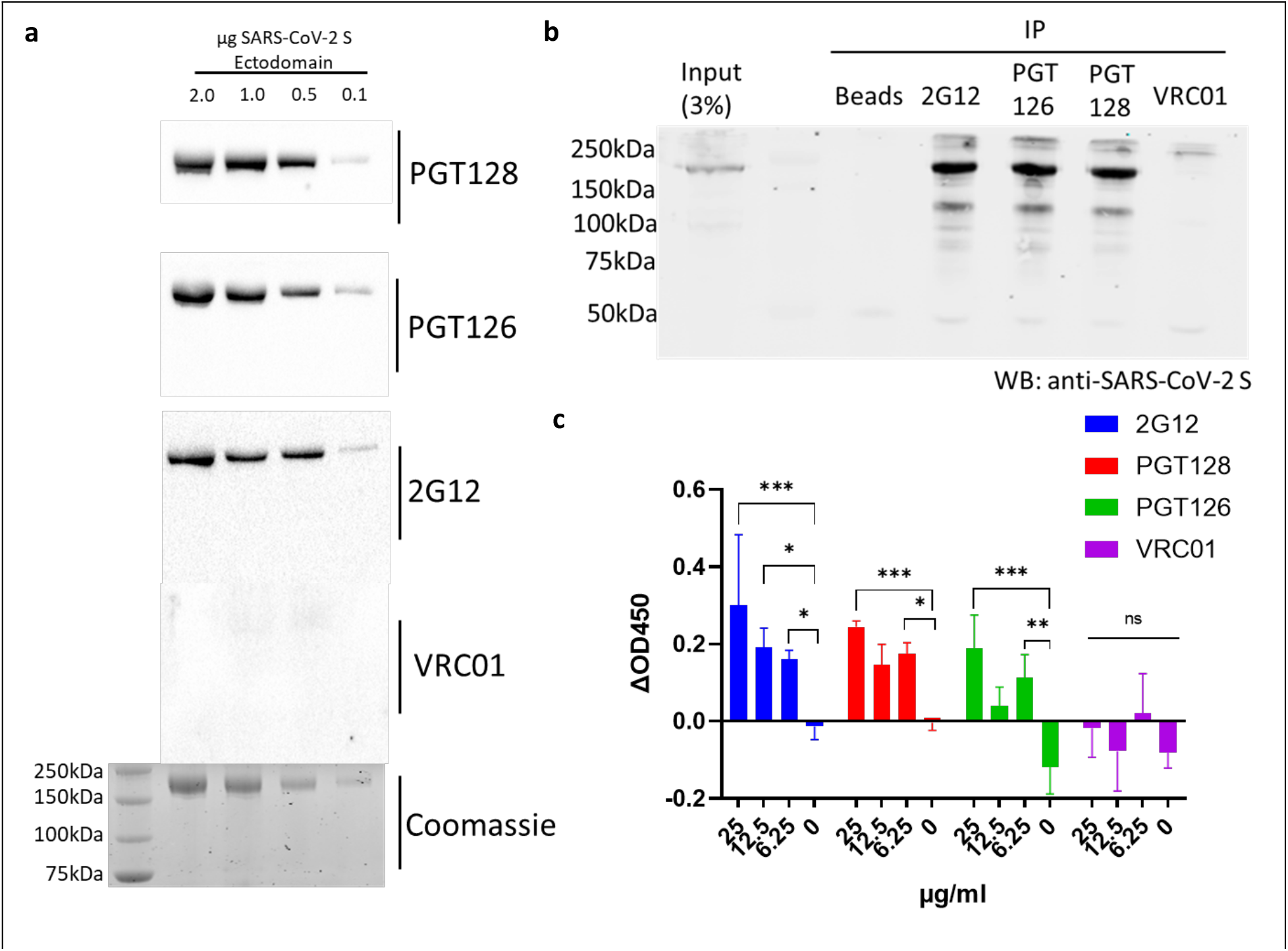
SARS-CoV-2 S binding capabilities of selected cross-reactive anti-gp120 antibodies. **(a)** Immunoreactivity of selected anti-gp120 antibodies with the SARS-CoV-2 S ectodomain was assessed via western blot, membranes were probed with the indicated antibodies prior to detection via HRP-anti-human IgG. A Coomassie-stained gel is included as a loading control. **(b)** Immunoprecipitation (IP) of full-length SARS-CoV-2 S by selected anti-gp120 antibodies. Lysates generated from HEK293T cells transiently expressing full-length SARS-CoV-2 S were incubated with the indicated antibodies and subjected to immunoprecipitation using protein A beads prior to Western blot (WB) analysis with a commercially available antibody targeting SARS-CoV-2 S. An immunoprecipitation condition using protein A beads alone is included as a control. Shown is a representative blot from 2 independent experiments. **(c)** Cell-based ELISA of cell-associated full-length SARS-CoV-2 S binding by the indicated anti-gp120 antibodies. Assays were carried out on chemically fixed HEK293T cells either transfected with plasmid encoding full-length SARS-CoV-2 S, or empty plasmid (mock). The difference in signal between these conditions is presented. Experiments were done in triplicate; error bars indicate standard deviation (n = 3). Statistical significance was tested by two-way ANOVA with Dunnett post-test (p > 0.05 [ns, not significant], p ≤ 0.05 [∗], p ≤ 0.01 [∗∗], p ≤ 0.001 [∗∗∗]).

Having characterized the cross-reactivities of 2G12, PGT128 and PGT126 with the SARS-CoV-2 spike, we proceeded to investigate the potential contribution of glycans in these interactions. We had observed abolished cross-reactivities when using casein-based blocking buffers (Figure S3) compared to BSA-based blocking buffers (Figure 1) in our initial cross-reactivity screens. Given the high carbohydrate content of casein(27), it suggested a possibility that these interactions may be glycan sensitive. Importantly, interactions between V_H_-FC ab8 and the SARS-CoV-2 spike ectodomain were not perturbed in casein-based buffer (Figure S2a-b). We first assessed these cross-reactive interactions in the presence of methyl α-d-mannopyranoside, a stabilized mannose analogue, via ELISA (Figure 4a). Disruption of cross-reactivity was observed for all three antibodies with increasing concentrations of methyl α-d-mannopyranoside, demonstrating the glycan sensitivity of these interactions. Expectedly, V_H_-FC ab8 displayed glycan insensitive binding in this assay, consistent with its RBD epitope within the SARS-CoV-2 spike(23). We next evaluated the cross-reactivities exhibited by these antibodies with differentially glycosylated SARS-CoV-2 spike preparations. We expressed SARS-CoV-2 S ectodomain in cells grown either in the presence or absence of kifunensine, a mannosidase inhibitor which facilitates the production of highly glycosylated, high mannose glycoproteins(28) (Figure S1). SDS-PAGE analysis of the resulting purified proteins revealed SARS-CoV-2 S ectodomain produced in kifunensine treated cells exhibits a hindered electrophoretic mobility relative to ectodomain produced in untreated cells (Figure 4b), consistent with the higher extent of glycosylation expected with kifunensine treatment. All three antibodies exhibited increased relative affinities and observed extents of binding with SARS-CoV-2 S ectodomain produced in cells treated with kifunensine compared to ectodomain produced in untreated cells (Figure 4c-d), further highlighting the participation of glycans within these cross-reactive interactions. As expected, V_H_-FC ab8 remained insensitive to the spike glycosylation state (Figure S2c). Taken together, these results implicate 2G12, PGT128 and PGT126 as targeting cross-reactive glyco-epitopes within the SARS-CoV-2 spike.

**Figure 4.**
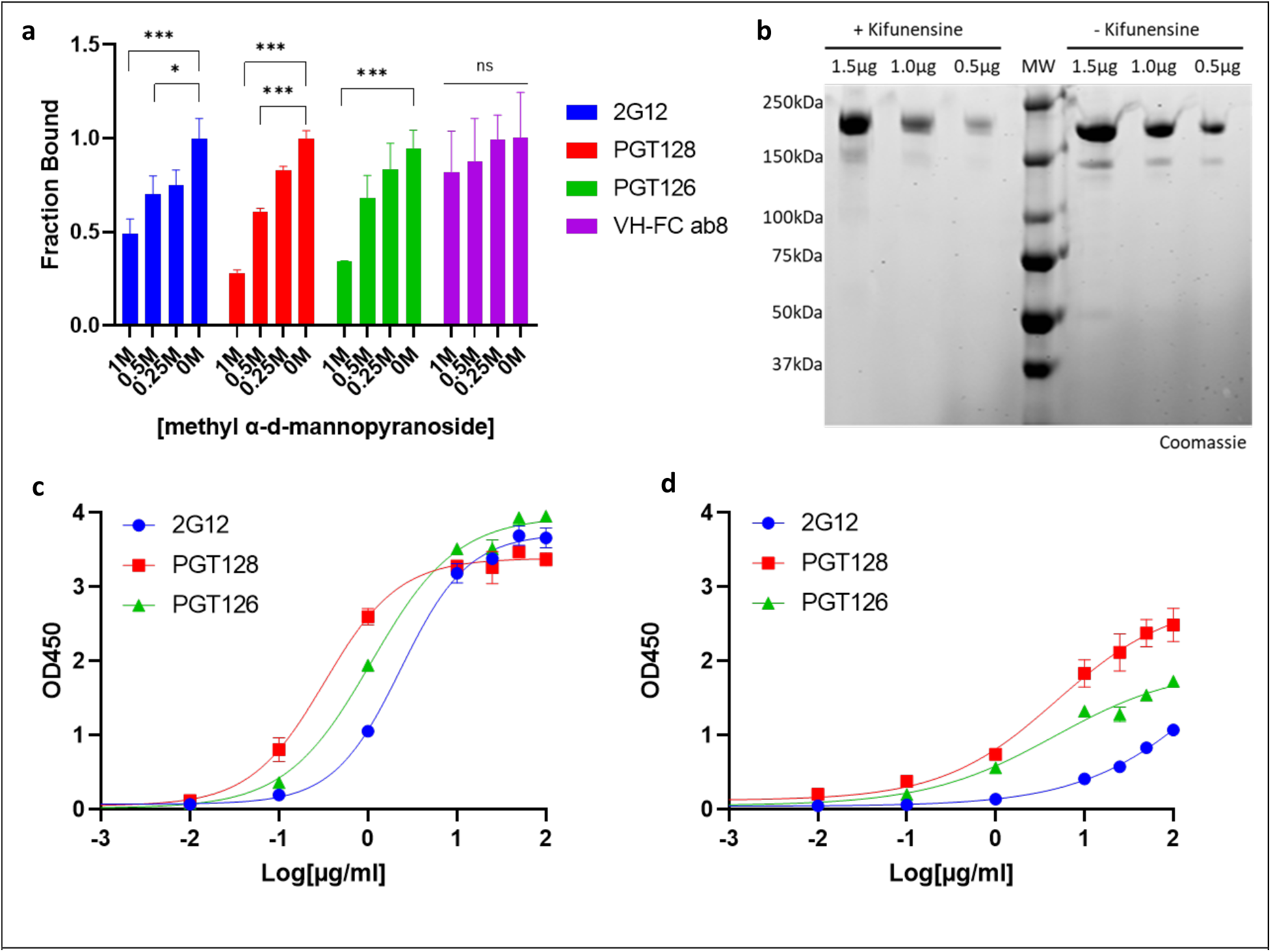
2G12, PGT128 and PGT126 cross-react with the SARS-CoV-2 Spike in a glycan dependent manner. **(a)** SARS-CoV-2 S ectodomain binding by anti-gp120 antibodies in the presence of methyl α-d-mannopyranoside. Plates were coated with SARS-CoV-2 S ectodomain and incubated with dilutions of methyl α-d-mannopyranoside along with a constant amount of the indicated antibodies. Antibody binding was quantified via ELISA. **(b)** SDS-PAGE analysis of varying amounts of purified SARS-CoV-2 S ectodomain expressed in cells either in the presence or absence of kifunensine. (MW: molecular weight ladder). **(c – d)** ELISA analysis of anti-gp120 antibody interactions with SARS-CoV-2 S ectodomain expressed in cells either in the presence **(c)** or absence of kifunensine **(d)**. Experiments were done in triplicate; error bars indicate standard deviation (n = 3). Statistical significance was tested by two-way ANOVA with Dunnett post-test (p > 0.05 [ns, not significant], p ≤ 0.05 [∗], p ≤ 0.01 [∗∗], p ≤ 0.001 [∗∗∗]).

## Discussion

We have identified a novel set of cross-reactive interactions between the SARS-CoV-2 spike and broadly neutralizing anti-HIV antibodies. A recent study(15) has demonstrated such cross-reactivity using Fab-dimerized antibodies such as 2G12, which target high mannose epitopes with gp120, and here we have extended these results to include anti-HIV antibodies which recognize gp120 at epitopes involving both peptide and glycans. Using a combination of immunoblotting, ELISA, and immunoprecipitation experiments, we demonstrate PGT128 and PGT126 bind both ectodomain and full-length SARS-CoV-2 S constructs, under denaturing and native conditions, in a glycan dependent manner. Using pseudo-typed viral entry assays, we demonstrate these cross-reactivities to be non-neutralizing, and likely incapable of mediating ADE via FcγRII receptor engagement.

In response to the global COVID-19 pandemic, a plethora of SARS-CoV-2 serological testing strategies have emerged(4–8), with the goal of providing epidemiological, diagnostic, and prognostic insight to health care providers. Such strategies have primarily focused on the detection of antibodies which bind the SARS-CoV-2 spike or nucleocapsid (N) proteins, however, recent studies have failed to demonstrate an appreciable predictive value between antibody titers and viral neutralization, disease status, and disease progression(16–19). While concurrent assessment of aspects of cellular immunity will likely contribute predictive value in such studies, efforts to further characterize humoral responses beyond serological detection of antigen-binding antibodies can provide a critical context for the interpretation of results. Assessments of aggregate antibody titers are incapable of differentiating between low affinity, non-neutralizing and high affinity, neutralizing antibodies, limiting their clinical utility. Cross-reactive antibodies have the potential to contribute confounding effects in such serological studies, and our results exemplify this possibility as we demonstrate the ability for low affinity, non-neutralizing, cross-reactive antibodies to bind the SARS-CoV-2 spike at antigenic glyco-epitopes. These results underscore the potential value of including neutralizing assays when evaluating patient antibody responses to SARS-CoV-2, and the importance of not relying solely on methods that only detect antibody-antigen interactions to evaluate humoral responses.

### Experimental Procedures

### Cell culture and DNA constructs

Expi293 cells were cultured in Expi293 expression medium, according to the manufacturer’s specifications. HEK293T (ATCC® CRL-3216) and HEK293T-ACE2 (BEI resources NR-52511) were cultured at 37°C, 5% CO_2_ in Dulbecco’s Modified Eagle medium (DMEM) supplemented with 10% fetal bovine serum (FBS) and 100 U/mL of penicillin–streptomycin. K-562 (ATCC® CCL-243) was cultured at 37°C, 5% CO_2_ in RPMI 1640 media supplemented with 10% FBS and 100 U/mL of penicillin–streptomycin. The codon-optimized SARS-CoV-2 2P S protein ectodomain construct (GenBank: <u>YP_009724390.1</u>) was C-terminally tagged with 8xHis and a twin Strep-tag and cloned into the mammalian expression vector pcDNA 3.1 (Synbio). The full-length SARS-CoV-2 S construct, pTwist-EF1alpha-SARS-CoV-2-S-2xStrep-IRES-Puro was a gift from Dr. Nevan Krogan.

#### Antibodies

All anti-gp120 IgG antibodies were obtained through the NIH AIDS Reagent Program, Division of AIDS, NIAID, NIH: Anti-HIV-1 gp120 Monoclonal (2G12) from Polymun Scientific, Anti-HIV-1 gp120 Monoclonal (PG9) from IAVI (cat# 12149), Anti-HIV-1 gp120 Monoclonal (PG16) from IAVI, Anti-HIV-1 gp120 Monoclonal (PGT121) from IAVI (cat# 12343), Anti-HIV-1 gp120 Monoclonal (PGT128) from IAVI (cat# 13352), Anti-HIV-1 gp120 Monoclonal (PGT145) from IAVI (cat# 12703), Cat# 12586 Anti-HIV-1 gp41/gp120 Monoclonal (35O22), from Drs. Jinghe Huang and Mark Connors, Anti-HIV-1 gp120 Monoclonal (10-1075) from Dr. Michel C. Nussenzweig (cat# 12477), Anti-HIV-1 gp120 Monoclonal (VRC01), from Dr. John Mascola (cat# 12033), Anti-HIV-1 gp120 Monoclonal (VRC03), from Dr. John Mascola (cat# 12032). VH-Fc ab8 was a gift from Dr. Dimiter S. Dimitrov.

#### Protein Expression and Purification

Transfections were performed as previously described (29, 30), with modifications. For SARS-CoV-2 2P S along with PGT126 and PGT128 Fab fragments, Expi293 cells were transfected at a density of 3 × 10^6^ cells/ml using linear polyethylenimine (PEI) (Polysciences). For Kifunensine treatment, cultures were treated with 5 µM Kifunensine 3 hrs post-transfection. At 24 hrs post-transfection, cultures were supplemented with 2.2 mM valproic acid. The supernatant was harvested by centrifugation after 5 days, filtered and loaded onto a 5 mL HisTrap Excel column (Cytiva). The column was washed with buffer (20 mM Tris pH 8.0, 500 mM NaCl, 20 mM imidazole) and the protein was eluted with buffer (20 mM Tris pH 8.0, 500 mM NaCl, 500 mM imidazole). Purified protein was concentrated and loaded onto a Superose 6 column (Cytiva) equilibrated with GF buffer (20 mM Tris pH 8.0 and 150 mM NaCl). Peak fractions were pooled, concentrated and flash frozen.

#### Enzyme-linked immunosorbent assay (ELISA)

100µl of SARS-CoV-2 2P S protein preparations were coated onto 96-well MaxiSorp™ plates at 1µg/ml in PBS overnight at 4°C. All washing steps were performed 5 times with PBS + 0.05% Tween 20 (PBS-T). After washing, wells were either incubated with BSA-based blocking buffer (PBS-T + 2% BSA) or Casein-based blocking buffer (TBS + 0.05% Tween 20 +1% Casein) for 1hr at room temperature. After washing, wells were incubated with dilutions of primary antibodies in either BSA-based blocking buffer or Casein-based blocking buffer for 2hrs at room temperature. After washing, wells were incubated with goat anti-human IgG (Jackson ImmunoResearch) at a 1:10,000 dilution in either BSA-based blocking buffer or Casein blocking buffer, for 1 hr at room temperature. After washing, the substrate solution (Pierce™ 1-Step™) was used for colour development according to the manufacturer’s specifications. Optical density at 450nm was read on a Varioskan Lux plate reader (Thermo Fisher Scientific). The same protocol was conducted for methyl α-d-mannopyranoside competition assays, keeping the concentration of the indicated primary antibodies at 5µg/ml while including dilutions of methyl α-d-mannopyranoside as indicated.

#### Cell-based ELISA

HEK293T cells were seeded in 96 well plates and transfected with either a plasmid encoding the full-length SARS-CoV-2 spike (pLVX-EF1alpha-SARS-CoV-2-S-2xStrep-IRES-Puro) or mock plasmid (pcDNA3.1) using branched PEI (Sigma). Media was switched 24hrs post-transfection. At 48hrs post-transfection, cells were washed 5 times with PBS prior to fixation with 4% paraformaldehyde in media for 30 minutes at 4°C. All further washing steps were performed 5 times with PBS + 0.05% Tween 20 (PBS-T). Cells were washed prior to blocking in blocking buffer (PBS-T + 2% BSA) for 1 hr at room temperature. After washing, cells were incubated with dilutions of primary antibodies in blocking buffer for 2 hrs at room temperature. After washing, cells were incubated with goat anti-human IgG (Jackson ImmunoResearch) at a 1:5,000 dilution in blocking buffer for 1 hr at room temperature. After washing, substrate solution (Pierce™ 1-Step™) was used for colour development according to the manufacturer’s specifications. Optical density at 450nm was read on a Varioskan Lux plate reader (Thermo Fisher Scientific). The difference in signals between cells transfected with full-length spike and mock plasmid was calculated.

#### Western Blotting

Purified SARS-CoV-2 2P spike ectodomain was subjected to SDS-PAGE and transferred onto nitrocellulose membranes prior to blocking in TBS + 0.05% Tween 20 (TBS-T) + 2% BSA for 1 hr at room temperature. Membranes were then incubated with the indicated anti-gp120 antibodies at 2µg/ml in TBS-T+ 2% BSA overnight at 4°C. After washing, membranes were incubated with goat anti-human IgG (Jackson ImmunoResearch) at a 1:5,000 dilution in TBS-T+ 2% BSA for 1 hr at room temperature. After washing, membranes were visualized using SuperSignal™ chemiluminescent substrate (Thermo Fisher Scientific).

### Immunoprecipitations

Immunoprecipitations were performed using the Dynabeads immunoprecipitation kit (Invitrogen). 100 µL of Dynabeads was incubated with 10 µg of either PGT126, PGT128, 2G12, VRC01, or PBS + 0.05% Tween 20 (PBS-T) as a beads-only control for 30 min at room temperature. Cellular lysates were generated from HEK-293T cells transiently expressing full-length SARS-CoV-2 spike (transfections as described for cell-based ELISA). Cells were solubilized in ice-cold 20 mM Tris-HCl, pH 7.4, 150 mM NaCl, 1 mM EDTA and 1% Triton X-100 with 1mM PMSF added fresh, and further disrupted via sonication. Lysates were then centrifuged at 12,000g for 2 mins and the supernatant was retrieved for immunoprecipitation input. After washing 3 times with 1 ml of PBS-T, antibody-bead complexes were incubated with lysate for 1 hr at room temperature on a rocking platform. Beads were washed 3 times with PBS-T and bound proteins were eluted directly into 4x Laemmli buffer with 10% dithiothreitol and boiled. The samples were then subjected to Western blot analysis and detected using an anti-SARS-CoV-2 spike glycoprotein antibody (Abcam – ab27504).

#### Pseudoviral entry assays

SARS-CoV-2 S pseudotyped retroviral particles were produced in HEK293T cells as described previously(24). Briefly, a 3^rd^ generation lentiviral packaging system was utilized in combination with plasmids encoding the full-length SARS-CoV-2 spike, along with a transfer plasmid encoding luciferase and GFP as a dual reporter gene. Pseudoviruses were harvested 60 hrs after transfection, filtered with 0.45 µm PES filters, and frozen. For cell-entry and neutralization assays, HEK293T-ACE2 cells were seeded in 96 well plates at 500,000 cells per well. The next day, pseudoviral preparations were incubated with dilutions of the indicated antibodies or media alone for 1 hr at 37°C prior to addition to cells and incubation for 48 hrs. Cells were then lysed and luciferase activity assessed using the ONE-Glo™ EX Luciferase Assay System (Promega) according to the manufacturer’s specifications. Detection of relative luciferase units was carried out using a Varioskan Lux plate reader (Thermo Fisher Scientific).

For ADE evaluation, human lymphoblast K562, which expresses FcγRII was utilized in the place of HEK293-T cells as described above. ADE was determined by comparing relative luciferase signals in the presence of the indicated cross-reactive antibodies to signals in the absence of these antibodies (pseudovirus only).

#### Statistical Analysis

One-way or two-way analysis of variance (ANOVA) with Dunnett or Sidak’s post-test was used to test for statistical significance. Only p values of 0.05 or lower were considered statistically significant (p > 0.05 [ns, not significant], p ≤ 0.05 [∗], p ≤ 0.01 [∗∗], p ≤ 0.001 [∗∗∗]). For all statistical analyses, GraphPad Prism 8 software package was used (GraphPad Software).

## Acknowledgments

We would like to thank Drs. Sagar Chittori and Shanti Swaroop Srivastava for helpful discussions and Steven Zhou for technical assistance with protein production. Work in the Subramaniam laboratory is supported by a Canada Excellence Research Chair Award and a grant from Genome BC, Canada. D.M. is supported by a CIHR Frederick Banting and Charles Best Canada Graduate Scholarship Master’s Award (CGS-M).

## Author contributions

D.M. and S.S. conceived the study. K.L. and D.M. performed protein purification. D.M conducted the experiments and data analysis. D.M., K.L., S.S., performed writing, review, and editing.

---

**Table S1.** HIV gp120 epitopes recognized by antibodies selected for SARS-CoV-2 S cross reactivity screening.

| Antibody | gp120 epitope | Reference |
| --- | --- | --- |
| 2G12 | N-linked glycans in the C2, C3, V4, and C4 domains | (31) |
| PGT121 | N332-centered oligomannose patch on the V3 loop, GDIR motif in V3 loop | (32–34) |
| PGT126 | N332-centered oligomannose patch of the V3 loop, GDIR motif in V3 loop | (32–34) |
| PGT128 | V3-glycan, GDIR motif in V3 loop | (32, 35, 36) |
| PGT145 | V1-V2 Glycans, residues within C-strand of V1/V2 and the C1 region near the base of V1 | (32, 37) |
| PG9 | V1-V2 Glycans,<br>residues within C<br>strand of V1/V2 | (38, 39) |
| PG16 | V1-V2 Glycans,<br>residues within V1–V2 | (38, 40, 41) |
| 10-1074 | V3-glycan, N332<br>glycosylation<br>dependent, GDIR<br>motif in V3 loop | (42, 43) |
| 35O22 | Glycan and peptides at<br>the gp120–gp41<br>interface | (44) |
| VRC01 | CD4-binding site | (45) |
| VRC03 | CD4-binding site | (45) |

**Figure S1.**
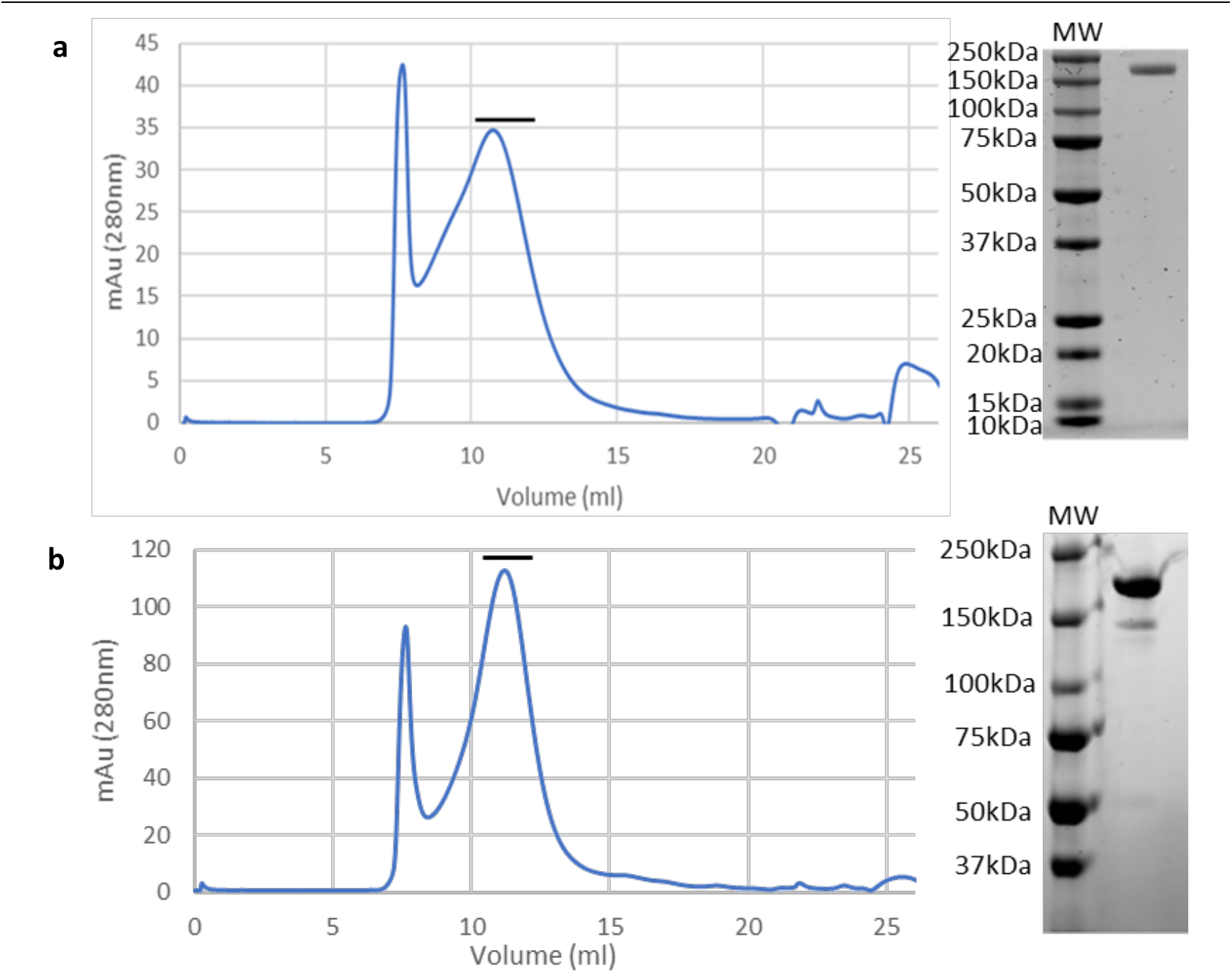
Purification of the SARS-CoV-2 S ectodomain expressed in cells either in the presence or absence of kifunensine. **(a**,**b)** Size exclusion chromatography profile of affinity purified SARS-CoV-2 S ectodomain expressed in cells either in the absence **(a)** or presence **(b)** of kifunensine run on a Superose 6 10/30 column, along with SDS-PAGE analyses of pooled and concentrated SARS-CoV-2 S ectodomain. The black line indicates fractions retrieved and pooled for SDS-PAGE analysis. Gels were stained with Coomassie. (MW: molecular weight ladder).

**Figure S2.**
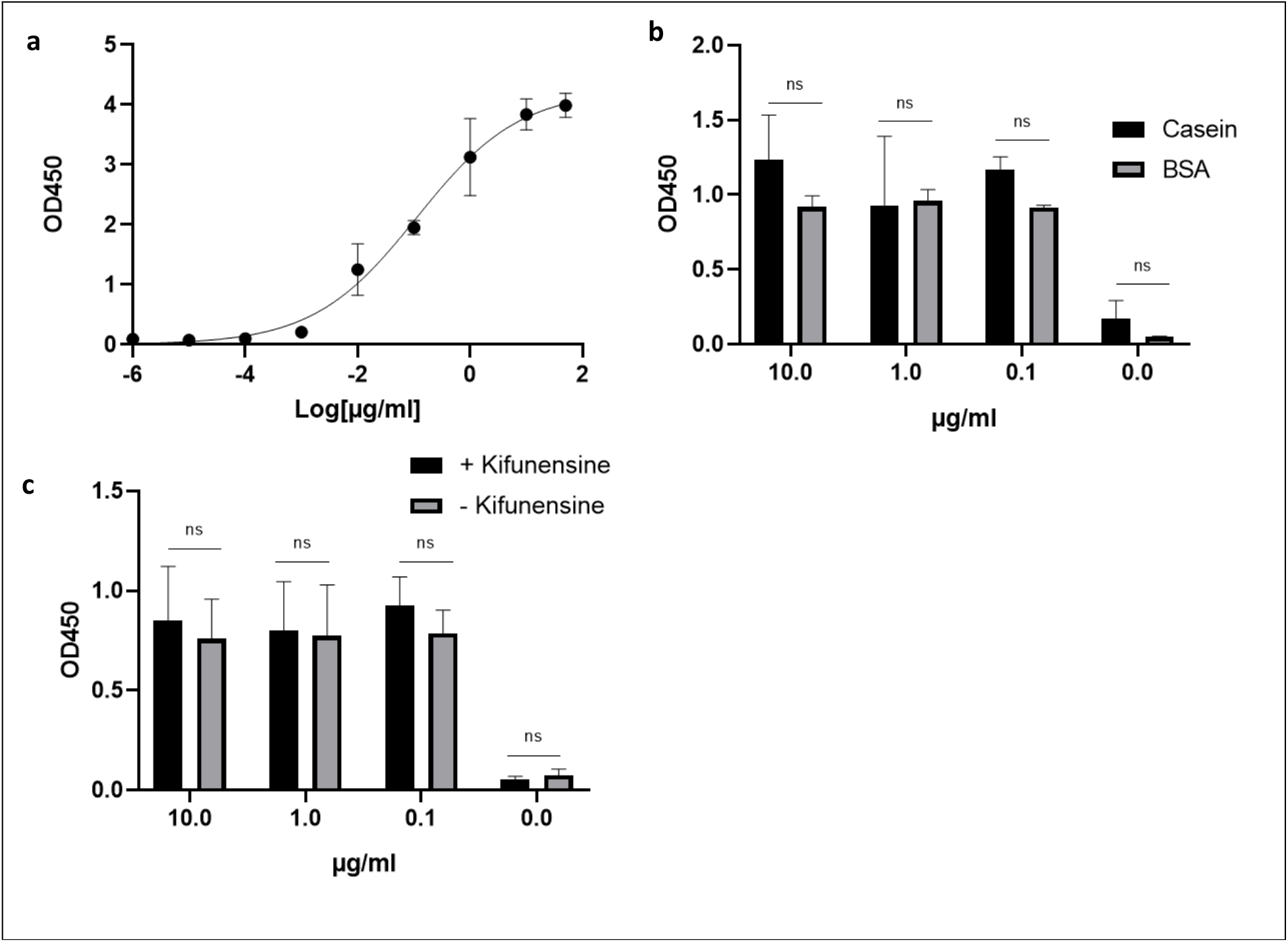
Analysis of VH-Fc ab8 – SARS-CoV-2 spike ectodomain interactions via ELISA. **(a)** SARS-CoV-2 S binding by VH-FC ab8, carried out in casein-based buffer. **(b)** SARS-CoV-2 S binding by VH-FC ab8 in the presence of BSA or casein-based buffers. **(c)** Interactions between VH-FC ab8 and SARS-CoV-2 S produced in cells grown in the presence or absence of kifunensine. Experiments were done in triplicate; error bars indicate standard deviation (n = 3). Statistical significance was tested by two-way ANOVA with Sidak’s post-test (p > 0.05 [ns, not significant], p ≤ 0.05 [∗], p ≤ 0.01 [∗∗], p ≤ 0.001 [∗∗∗]).

**Figure S3.**
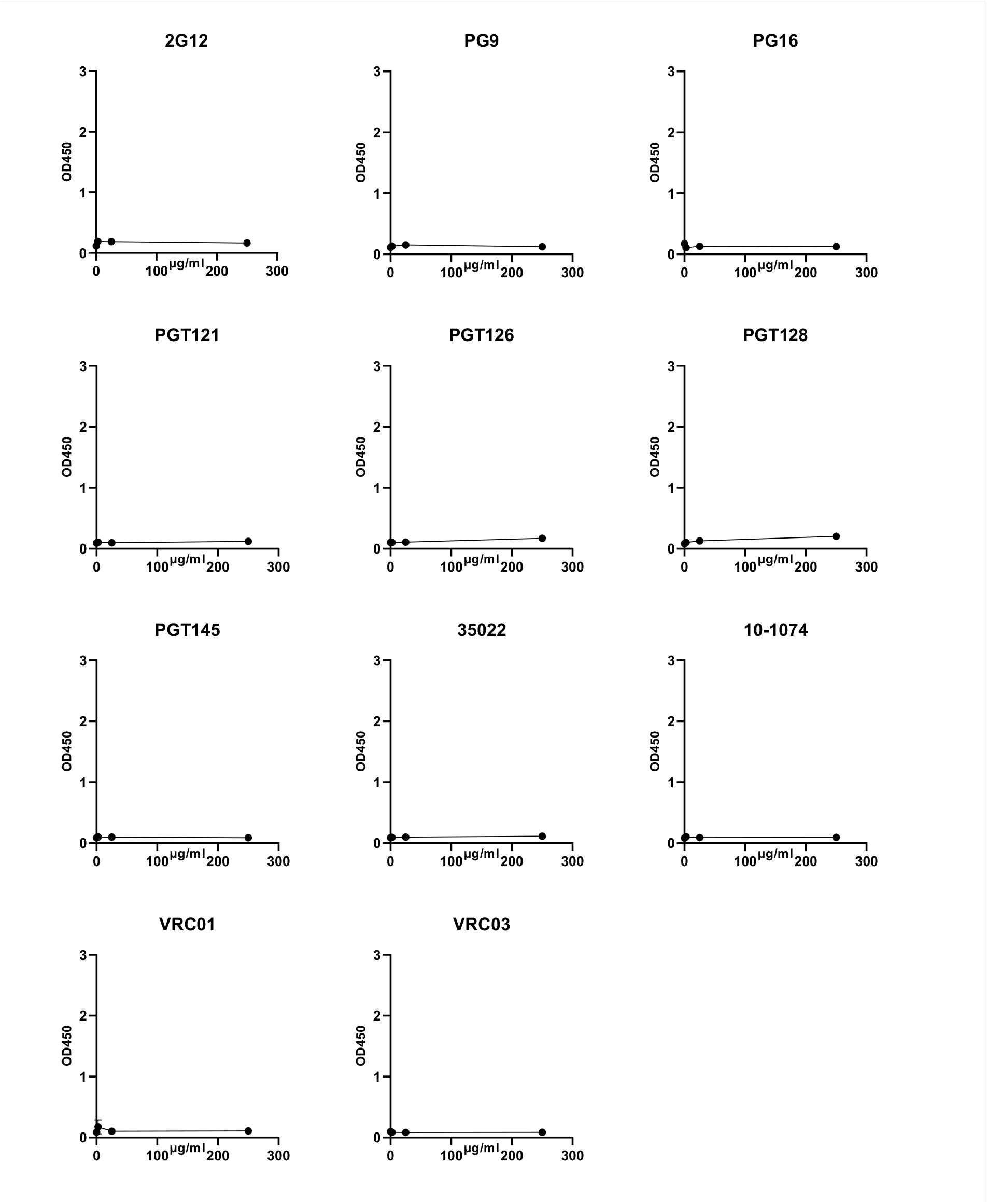
ELISA screen of glycan directed anti-gp120 antibody cross-reactivities to the SARS-CoV-2 Spike using the casein-based buffer. Serial dilutions of the indicated mAbs were assessed for SARS-CoV-2 S protein binding. VRC01 and VRC03 target the CD4 binding site within gp120 and are included here as negative controls. All ELISAs were performed using casein-based buffers (see methods). Experiments were done in duplicate; error bars indicate standard deviation (n = 2).

## References

1. Gallagher, T. M., and Buchmeier, M. J. (2001) Coronavirus spike proteins in viral entry and pathogenesis. Virology. 279, 371–374

2. Walls, A. C., Park, Y.-J., Tortorici, M. A., Wall, A., McGuire, A. T., and Veesler, D. (2020) Structure, Function, and Antigenicity of the SARS-CoV-2 Spike Glycoprotein. Cell. 181, 281-292.e6

3. Cai, Y., Zhang, J., Xiao, T., Peng, H., Sterling, S. M., Walsh, R. M., Rawson, S., Rits-Volloch, S., and Chen, B. (2020) Distinct conformational states of SARS-CoV-2 spike protein. Science (80-.). 369, 1586–1592

4. Wölfel, R., Corman, V. M., Guggemos, W., Seilmaier, M., Zange, S., Müller, M. A., Niemeyer, D., Jones, T. C., Vollmar, P., Rothe, C., Hoelscher, M., Bleicker, T., Brünink, S., Schneider, J., Ehmann, R., Zwirglmaier, K., Drosten, C., and Wendtner, C. (2020) Virological assessment of hospitalized patients with COVID-2019. Nature. 581, 465–469

5. Kohmer, N., Westhaus, S., Rühl, C., Ciesek, S., and Rabenau, H. F. (2020) Brief clinical evaluation of six high-throughput SARS-CoV-2 IgG antibody assays. J. Clin. Virol. 129, 104480

6. Charlton, C. L., Kanji, J. N., Johal, K., Bailey, A., Plitt, S. S., MacDonald, C., Kunst, A., Buss, E., Burnes, L. E., Fonseca, K., Berenger, B. M., Schnabl, K., Hu, J., Stokes, W., Zelyas, N., and Tipples, G. (2020) Evaluation of Six Commercial Mid-to High-Volume Antibody and Six Point-of-Care Lateral Flow Assays for Detection of SARS-CoV-2 Antibodies. J. Clin. Microbiol. 58, e01361–20

7. Amanat, F., Stadlbauer, D., Strohmeier, S., Nguyen, T. H. O., Chromikova, V., Mcmahon, M., Jiang, K., Arunkumar, G. A., Jurczyszak, D., Polanco, J., Bermudez-gonzalez, M., Kleiner, G., Aydillo, T., Miorin, L., Fierer, D. S., Lugo, L. A., Kojic, E. M., Stoever, J., Liu, S. T. H., Cunningham-rundles, C., Felgner, P. L., Moran, T., García-sastre, A., Caplivski, D., Cheng, A. C., and Kedzierska, K. (2020) A serological assay to detect SARS-CoV-2 seroconversion in humans. Nat. Med. 10.1038/s41591-020-0913-5

8. Montesinos, I., Gruson, D., Kabamba, B., Dahma, H., Van den Wijngaert, S., Reza, S., Carbone, V., Vandenberg, O., Gulbis, B., Wolff, F., and Rodriguez-Villalobos, H. (2020) Evaluation of two automated and three rapid lateral flow immunoassays for the detection of anti-SARS-CoV-2 antibodies. J. Clin. Virol. 128, 104413

9. Zhu, Y., Yu, D., Han, Y., Yan, H., Chong, H., Ren, L., Wang, J., Li, T., and He, Y. (2020) Cross-reactive neutralization of SARS-CoV-2 by serum antibodies from recovered SARS patients and immunized animals. Sci. Adv. 10.1126/sciadv.abc9999

10. Wang, C., Li, W., Drabek, D., Okba, N. M. A., Haperen, R. Van Osterhaus, A.D M.E., Kuppeveld, F. J. M. Van, Haagmans, B. L., Grosveld, F., and Bosch, B. A human monoclonal antibody blocking SARS-CoV-2 infection. Nat. Commun. 10.1038/s41467-020-16256-y

11. Pinto, D., Park, Y., Beltramello, M., Walls, A. C., Tortorici, M. A., Bianchi, S., Jaconi, S., Culap, K., Zatta, F., Marco, A. De Peter, A., Guarino, B., Spreafico, R., Cameroni, E., Case, J. B., Chen, R. E., Havenar-daughton, C., Snell, G., Telenti, A., Virgin, H. W., Lanzavecchia, A., Diamond, M. S., Fink, K., Veesler, D., and Corti, D. (2020) Cross-neutralization of SARS-CoV-2 by a human monoclonal SARS-CoV antibody. Nature. 10.1038/s41586-020-2349-y

12. Ng, K. W., Faulkner, N., Cornish, G. H., Rosa, A., Harvey, R., Hussain, S., Ulferts, R., Earl, C., Wrobel, A. G., Benton, D. J., Roustan, C., Bolland, W., Thompson, R., Agua-Doce, A., Hobson, P., Heaney, J., Rickman, H., Paraskevopoulou, S., Houlihan, C. F., Thomson, K., Sanchez, E., Shin, G. Y., Spyer, M. J., Joshi, D., textquoterightReilly, N., Walker, P. A., Kjaer, S., Riddell, A., Moore, C., Jebson, B. R., Wilkinson, M., Marshall, L. R., Rosser, E. C., Radziszewska, A., Peckham, H., Ciurtin, C., Wedderburn, L. R., Beale, R., Swanton, C., Gandhi, S., Stockinger, B., McCauley, J., Gamblin, S. J., McCoy, L. E., Cherepanov, P., Nastouli, E., and Kassiotis, G. (2020) Preexisting and de novo humoral immunity to SARS-CoV-2 in humans. Science (80-.). 10.1126/science.abe1107

13. Lustig, Y., Keler, S., Kolodny, R., Ben-Tal, N., Atias-Varon, D., Shlush, E., Gerlic, M., Munitz, A., Doolman, R., Asraf, K., Shlush, L. I., and Vivante, A. (2020) Potential Antigenic Cross-reactivity Between Severe Acute Respiratory Syndrome Coronavirus 2 (SARS-CoV-2) and Dengue Viruses. Clin. Infect. Dis. 10.1093/cid/ciaa1207

14. Nath, H., Mallick, A., Roy, S., Sukla, S., Basu, K., De, A., and Biswas, S. (2020) Dengue antibodies can cross-react with SARS-CoV-2 and vice versa-Antibody detection kits can give false-positive results for both viruses in regions where both COVID-19 and Dengue co-exist. 10.1101/2020.07.03.20145797

15. [preprint] Acharya, P., Williams, W., Henderson, R., Janowska, K., Manne, K., Parks, R., Deyton, M., Sprenz, J., Stalls, V., Kopp, M., Mansouri, K., Edwards, R. J., Meyerhoff, R. R., Oguin, T., Sempowski, G., Saunders, K., and Haynes, B. F. (2020) A glycan cluster on the SARS-CoV-2 spike ectodomain is recognized by Fab-dimerized glycan-reactive antibodies. bioRxiv Prepr. Serv. Biol. 10.1101/2020.06.30.178897

16. Ding, S., Laumaea, A., Benlarbi, M., Beaudoin-bussi, G., Gasser, R., Medjahed, H., Pancera, M., Stamatatos, L., and Mcguire, A. T. (2020) Antibody Binding to SARS-CoV-2 S Glycoprotein Correlates with but Does Not Predict Neutralization. 2, 1–9

17. Escribano, P., Uría, A.Á., Alonso, R., Catalán, P., and Alcalá, L. (2020) Detection of SARS - CoV - 2 antibodies is insufficient for the diagnosis of active or cured COVID - 19. Sci. Rep. 10.1038/s41598-020-76914-5

18. Phipps, W. S., SoRelle, J. A., Li, Q.-Z., Mahimainathan, L., Araj, E., Markantonis, J., Lacelle, C., Balani, J., Parikh, H., Solow, E. B., Karp, D. R., Sarode, R., and Muthukumar, (2020) SARS-CoV-2 Antibody Responses Do Not Predict COVID-19 Disease Severity. Am. J. Clin. Pathol. 154, 459–465

19. Criscuolo, E., Diotti, R. A., Strollo, M., Rolla, S., Ambrosi, A., Locatelli, M., Burioni, R., Mancini, N., Clementi, M., and Clementi, N. Weak correlation between antibody titers and neutralizing activity in sera from SARS-CoV-2 infected subjects. J. Med. Virol. https://doi.org/10.1002/jmv.26605

20. Bagdonaite, I., and Wandall, H. H. (2018) Global aspects of viral glycosylation. Glycobiology. 28, 443–467

21. Doores, K. J. (2015) The HIV glycan shield as a target for broadly neutralizing antibodies. FEBS J. 282, 4679–4691

22. Watanabe, Y., Allen, J. D., Wrapp, D., McLellan, J. S., and Crispin, M. (2020) Site-specific glycan analysis of the SARS-CoV-2 spike. Science (80-.). 369, 330 LP – 333

23. Li, W., Scha, A., Kulkarni, S. S., Subramaniam, S., Baric, R. S., Dimitrov, D. S., Kulkarni, S. S., Liu, X., Martinez, D. R., Chen, C., and Sun, Z. (2020) Article High Potency of a Bivalent Human V H Domain in SARS-CoV-2 Animal Models ll Article High Potency of a Bivalent Human V H Domain in SARS-CoV-2 Animal Models. Cell. 10.1016/j.cell.2020.09.007

24. Crawford, K. H. D., Eguia, R., Dingens, A. S., Loes, A. N., Malone, K. D., Wolf, C. R., Chu, H. Y., Tortorici, M. A., Veesler, D., Murphy, M., Pettie, D., King, N. P., Balazs, A. B., and Bloom, J. D. (2020) Protocol and Reagents for Pseudotyping Lentiviral Particles with SARS-CoV-2 Spike Protein for Neutralization Assays

25. [preprint] Wu, F., Yan, R., Liu, M., Liu, Z., Wang, Y., Luan, D., Wu, K., Song, Z., Sun, T., Ma, Y., Zhang, Y., Wang, Q., Li, X., Ji, P., Li, Y., Li, C., Wu, Y., Ying, T., Wen, Y., Jiang, S., Zhu, T., Lu, L., Zhang, Y., Zhou, Q., and Huang, J. (2020) Antibody-dependent enhancement (ADE) of SARS-CoV-2 infection in recovered COVID-19 patients: studies based on cellular and structural biology analysis. medRxiv. 10.1101/2020.10.08.20209114

26. Chiofalo, M. S., Teti, G., Goust, J. M., Trifiletti, R., and La Via, M. F. (1988) Subclass specificity of the Fc receptor for human IgG on K562. Cell. Immunol. 114, 272–281

27. Reynolds, L. M., Henneberry, G. O., and Baker, B. E. (1959) Studies on Casein. II. The Carbohydrate Moiety of Casein. J. Dairy Sci. 42, 1463–1471

28. Tropea, E. (1990) Kifunensine, a Potent Inhibitor Mannosidase I * of the Glycoprotein Processing. J. Biol. Chem. 265, 15599–15605

29. Portolano, N., Watson, P. J., Fairall, L., Millard, C. J., Milano, C. P., Song, Y., and Cowley, S. M. (2014) Recombinant Protein Expression for Structural Biology in HEK 293F Suspension Cells?: A Novel and Accessible Approach 2. Protein-complex Purification from Whole Cell Extract. 1, 1–8

30. Vazquez-lombardi, R., Nevoltris, D., Luthra, A., Schofield, P., Zimmermann, C., and Christ, D. (2017) Transient expression of human antibodies in mammalian cells. Nat. Publ. Gr. 13, 99–117

31. Trkola, A., Purtscher, M., Muster, T., Ballaun, C., Buchacher, A., Sullivan, N., Srinivasan, K., Sodroski, J., Moore, J. P., and Katinger, H. (1996) Human monoclonal antibody 2G12 defines a distinctive neutralization epitope on the gp120 glycoprotein of human immunodeficiency virus type 1. J. Virol. 70, 1100–1108

32. Walker, L. M., Huber, M., Doores, K. J., Falkowska, E., Pejchal, R., Julien, J.-P., Wang, S.-K., Ramos, A., Chan-Hui, P.-Y., Moyle, M., Mitcham, J. L., Hammond, P. W., Olsen, O. A., Phung, P., Fling, S., Wong, C.-H., Phogat, S., Wrin, T., Simek, M. D., Investigators, P. G. P., Koff, W. C., Wilson, I. A., Burton, D. R., and Poignard, P. (2011) Broad neutralization coverage of HIV by multiple highly potent antibodies. Nature. 477, 466–470

33. Julien, J., Sok, D., Khayat, R., Lee, J. H., Doores, K. J., Walker, L. M., Ramos, A., Diwanji, D. C., Pejchal, R., Cupo, A., Katpally, U., Depetris, R. S., Stanfield, R. L., Mcbride, R., Marozsan, A. J., Paulson, J. C., Sanders, R. W., Moore, J. P., Burton, D. R., Poignard, P., Ward, A. B., and Wilson, I. A. (2013) Broadly Neutralizing Antibody PGT121 Allosterically Modulates CD4 Binding via Recognition of the HIV-1 gp120 V3 Base and Multiple Surrounding Glycans. 10.1371/journal.ppat.1003342

34. Garces, F., Lee, J. H., Val, N. De Sanders, R. W., Ward, A. B., Wilson, I. A., Garces, F., Lee, J. H., Val, N., De Torrents, A., Pena, D., and Kong, L. (2015) Affinity Maturation of a Potent Family of HIV Antibodies Is Primarily Focused on Accommodating or Avoiding Glycans Article Affinity Maturation of a Potent Family of HIV Antibodies Is Primarily Focused on Accommodating or Avoiding Glycans. Immunity. 43, 1053–1063

35. Pejchal, R., Doores, K. J., Walker, L. M., Khayat, R., Huang, P.-S., Wang, S.-K., Stanfield, R. L., Julien, J.-P., Ramos, A., Crispin, M., Depetris, R., Katpally, U., Marozsan, A., Cupo, A., Maloveste, S., Liu, Y., McBride, R., Ito, Y., Sanders, R. W., Ogohara, C., Paulson, J. C., Feizi, T., Scanlan, C. N., Wong, C.-H., Moore, J. P., Olson, W. C., Ward, A. B., Poignard, P., Schief, W. R., Burton, D. R., and Wilson, I. A. (2011) A potent and broad neutralizing antibody recognizes and penetrates the HIV glycan shield. Science. 334, 1097–1103

36. Lee, J. H., Val, N., De Ward, A. B., Lee, J. H., Val, N., De Lyumkis, D., and Ward, A. B. (2015) Model Building and Refinement of a Natively Glycosylated HIV-1 Env Protein by High-Resolution Cryoelectron Microscopy Article Model Building and Refinement of a Natively Glycosylated HIV-1 Env Protein by High-Resolution Cryoelectron Microscopy. Struct. Des. 23, 1943–1951

37. Lee, J. H., Andrabi, R., Su, C., Wilson, I. A., Burton, D. R., and Ward, A. B. (2017) A Broadly Neutralizing Antibody Targets the Dynamic HIV Envelope Trimer Apex via a Long, Rigidified, and Anionic b -Hairpin Structure. Immunity. 46, 690–702

38. Walker, L. M., Phogat, S. K., Chan-Hui, P.-Y., Wagner, D., Phung, P., Goss, J. L., Wrin, T., Simek, M. D., Fling, S., Mitcham, J. L., Lehrman, J. K., Priddy, F. H., Olsen, O. A., Frey, S. M., Hammond, P. W., Investigators, P. G. P., Kaminsky, S., Zamb, T., Moyle, M., Koff, W. C., Poignard, P., and Burton, D. R. (2009) Broad and potent neutralizing antibodies from an African donor reveal a new HIV-1 vaccine target. Science. 326, 285–289

39. Mclellan, J. S., Pancera, M., Carrico, C., Gorman, J., Julien, J., Khayat, R., Louder, R., Pejchal, R., Sastry, M., Dai, K., Dell, S. O., Patel, N., Shahzad-ul-hussan, S., Yang, Y., Zhang, B., Zhou, T., Zhu, J., Boyington, J. C., Chuang, G., Diwanji, D., Georgiev, I., Kwon, Y., Do Lee, D., Louder, M. K., Moquin, S., Schmidt, S. D., Yang, Z., Bonsignori, M., Crump, J. A., Kapiga, S. H., Sam, N. E., Haynes, B. F., Burton, D. R., Koff, W. C., Walker, L. M., Phogat, S., Wyatt, R., Orwenyo, J., Wang, L., Arthos, J., Bewley, C. A., and Mascola, J. R. (2011) Structure of HIV-1 gp120 V1/V2 domain with broadly neutralizing antibody PG9. Nature. 10.1038/nature10696

40. Pancera, M., Shahzad-ul-hussan, S., Doria-rose, N. A., Mclellan, J. S., Bailer, R. T., Dai, K., Loesgen, S., Louder, M. K., Staupe, R. P., Yang, Y., Zhang, B., Parks, R., Eudailey, J., Lloyd, K. E., Blinn, J., Alam, S. M., Haynes, B. F., Amin, M. N., Wang, L., Burton, D. R., Koff, W. C., Nabel, G. J., Mascola, J. R., Bewley, C. A., and Kwong, P. D. (2013) Structural basis for diverse N-glycan recognition by HIV-1 – neutralizing V1 – V2 – directed antibody PG16. Nat. Publ. Gr. 10.1038/nsmb.2600

41. Pan, J., Peng, H., Chen, B., and Harrison, S. C. (2020) Cryo-EM Structure of Full-length HIV-1 Env Bound With the Fab of Antibody PG16. J. Mol. Biol. 432, 1158–1168

42. Shingai, M., Nishimura, Y., Klein, F., Mouquet, H., Donau, O. K., Plishka, R., Buckler-White, A., Seaman, M., Piatak Jr, M., Lifson, J. D., Dimitrov, D. S., Nussenzweig, M. C., and Martin, M. A. (2013) Antibody-mediated immunotherapy of macaques chronically infected with SHIV suppresses viraemia. Nature. 503, 277–280

43. Gristick, H. B., von Boehmer, L., West Jr, A. P., Schamber, M., Gazumyan, A., Golijanin, J., Seaman, M. S., Fätkenheuer, G., Klein, F., Nussenzweig, M. C., and Bjorkman, P. J. (2016) Natively glycosylated HIV-1 Env structure reveals new mode for antibody recognition of the CD4-binding site. Nat. Struct. Mol. Biol. 23, 906–915

44. Huang, J., Kang, B. H., Pancera, M., Lee, J. H., Tong, T., Feng, Y., Imamichi, H., Georgiev, I. S., Chuang, G.-Y., Druz, A., Doria-Rose, N. A., Laub, L., Sliepen, K., van Gils, M. J., de la Peña, A. T., Derking, R., Klasse, P.-J., Migueles, S. A., Bailer, R. T., Alam, M., Pugach, P., Haynes, B. F., Wyatt, R. T., Sanders, R. W., Binley, J. M., Ward, B., Mascola, J. R., Kwong, P. D., and Connors, M. (2014) Broad and potent HIV-1 neutralization by a human antibody that binds the gp41-gp120 interface. Nature. 515, 138–142

45. Wu, X., Yang, Z.-Y., Li, Y., Hogerkorp, C.-M., Schief, W. R., Seaman, M. S., Zhou, T., Schmidt, S. D., Wu, L., Xu, L., Longo, N. S., McKee, K., O’Dell, S., Louder, M. K., Wycuff, D. L., Feng, Y., Nason, M., Doria-Rose, N., Connors, M., Kwong, P. D., Roederer, M., Wyatt, R. T., Nabel, G. J., and Mascola, J. R. (2010) Rational design of envelope identifies broadly neutralizing human monoclonal antibodies to HIV-1. Science. 329, 856–861

